# IlvY is an important regulator of *Shigella* infection *in vitro* and *in vivo*

**DOI:** 10.1101/2020.07.28.220327

**Authors:** Mayumi K. Holly, Mark C. Anderson, Lesley M. Rabago, Azadeh Saffarian, Benoit S. Marteyn, Samuel L.M. Arnold

## Abstract

Shigellosis results from oral ingestion of the Gram-negative bacteria *Shigella*, and symptoms include severe diarrhea and dysentery. In the absence of vaccines, small molecule antibacterial drugs have provided treatment options for shigellosis. However, *Shigella* drug resistance is rapidly emerging, and *Shigella* strains with resistance to both third-generation cephalosporins and azithromycin have been identified in Asia. A re-conceptualization is needed regarding the development of therapeutics that target bacterial pathogens in order to reduce resistance development and alteration of gut microbiota, which is depleted upon treatment with wide spectrum antibiotics, thereby increasing susceptibility to subsequent enteric infections. A more organism-specific approach is to develop agents targeting virulence factors such as toxins, adhesins, invasins, quorum sensing, and protein secretion systems. For *Shigella*, there is interest in targeting transcription factors essential for *Shigella* infection *in vivo* rather than specific effectors. Here we describe the importance of the *Shigella* transcription factor IlvY in *Shigella* virulence *in vitro* and *in vivo*. This work included the development of a novel, oral mouse model of *Shigella* infection with wild-type adult mice. Unlike previous models, mice do not require antibiotic pretreatment or diet modifications. This mouse model was used to demonstrate the importance of IlvY for *Shigella in vivo* survival and that deletion of *ilvY* impacts host responses to infection. These results illustrate that IlvY is a potential therapeutic target for the treatment of shigellosis. In addition, the novel mouse model provides an exciting new opportunity to investigate therapeutic efficacy against *Shigella* infection and host responses to infection.

## Introduction

*Shigella* spp. are a Gram-negative, facultative enteric bacteria that are the causative agents of shigellosis [1]. Globally, *Shigella* causes 75 million cases each year annually and is the leading cause of childhood morbidity and mortality in the developing world [2]. This disease is characterized by watery, mucoid or bloody diarrhea and in more severe cases, fever and tenesmus [1]. Treatment of shigellosis is becoming more difficult due to an alarming increase in antimicrobial resistance (AMR) to World Health Organization recommended first- and second-line antibacterial therapeutics: ciprofloxacin and azithromycin, respectively. To compound this issue, no new classes of antibacterial therapeutics have been approved for clinical use to treat Gram-negative bacteria [3]. The Centers for Disease Control and Prevention has classified multidrug non-susceptible *Shigella* as a serious threat, and there is an urgent need of new therapeutics for treating shigellosis [4].

AMR has been emerging since the advent of antibiotics in the early 20^th^ century [3]. Traditionally antibacterial development has focused on targeting essential bacterial processes that are distinct from mammalian cellular processes. The rapid emergence of AMR following release into the clinic has largely been fueled by three factors: 1) misuse of antibacterial therapeutics for incorrect applications, 2) lack of completion of antibacterial course, and 3) targeting bacterial processes that are essential for bacterial survival in all environments [5]. The first two factors can be addressed at the level of the clinic, but the third factor represents an issue with the development of antibacterial therapeutics. By targeting and inhibiting essential bacterial processes, traditional antibiotics place significant selective pressure on bacteria to develop resistance to the treatment. Additionally, this resistance can be passed among different species of bacteria through horizontal gene transfer. Therefore, it is imperative to identify new therapeutic targets for treatment of bacterial infections.

Another avenue of therapeutic targets being explored focuses on bacteria transcription factors. Transcription factors as therapeutic targets have long been thought of as undruggable due to the difficulty of targeting the transcription factor - DNA interface. However, recent studies provide support for the idea of drugging transcription factors [6–9]. Transcription factors are an ideal target because they control gene expression programs that are important for infection and pathogenesis, while not being essential for bacterial cell survival. *Shigella* spp. encode a wide variety of different transcriptional programs that allow for effective execution of the different stages of *Shigella* infection and pathogenesis.

Infection of the colon by *Shigella* is a multi-stage process involving competition with the microbiome, survival in anaerobic and hypoxic environments, invasion of epithelial cells, direct cell-to-cell spread, and invasion of immune cells [1, 10]. Each stage is controlled by various transcription factors that turn on or off specific suites of genes. The master regulator of virulence gene expression is VirF, an AraC-like transcription factor[11]. *virF* expression is repressed by HN-S at 30°C, but upon transition to 37°C HN-S repression of *virF* is relieved allowing for synthesis of VirF. Given that VirF is the master regulator of virulence factor expression, it is currently being pursued as a drug target [11, 12]. However, in addition to VirF, it is likely there are additional *Shigella* transcription factors that remain to be investigated as therapeutic targets.

A *S. sonnei* transposon mutant library *in vivo* screen conducted in guinea pigs identified the transcription factor IlvY as essential for *S. sonnei* survival in vivo (Personal communication: Dr. Mark Anderson). IlvY regulates expression of *ilvC* which encodes IlvC, a ketol-acid reductoisomerase that catalyzes the conversion of acetolactate and acetohydroxybutyrate into products for use in the biosynthesis of two branched chain amino acids (BCAA) valine and isoleucine [13]. IlvY and BCAAs potentially play several roles during *Shigella* infection including response to nitric oxide (NO), amino acid starvation, and activation of virulence factors [14–16]. Due to its diverse roles, IlvY is a prime therapeutic target. We investigated the role of IlvY during *S. flexneri* and *S. sonnei in vitro* infection to determine whether IlvY is a suitable therapeutic target. Due to the lack of a robust mouse model for years, it has been difficult to translate findings *in vitro* to *in vivo* infection and pathogenesis. While there are existing mouse models for *Shigella* infection that use non-oral routes of infection [17–19], oral infection models require antibiotic pretreatment, diet modifications, and/or immunodeficient adult mice [20, 21]. For the first time, we describe an oral adult mouse model of *Shigella* infection that uses wild-type mice and does not require antibiotic pretreatment or diet modifications. Using this mouse model, we report that *Shigella* deficient in IlvY are attenuated for infection compared to wild-type *Shigella*. Our mouse model of *Shigella* now allows for translation of *in vitro* results to *in vivo* effects.

## Results

### IlvY is important for replication under nutrient limiting conditions

Ideally, transcription factors for therapeutic development are non-essential, involved in infection, and conserved across many pathogenic bacteria. Data from a previously performed *in vivo* transposon mutant screen suggested that the transcription factor IlvY was important for *Shigella in vivo* survival (Personal communication: Dr. Mark Anderson). In agreement with ideal criteria for transcription factors as therapeutic targets, IlvY is highly conserved within Enterobacterales (Fig. 1). IlvY regulates expression of two genes, itself and *ilvC*, which encodes a ketol-acid reductoisomerase [13, 22]. IlvC is part of both the valine and isoleucine biosynthetic pathways and functions to convert acetohydroxybutyrate or acetolactate to 2,3-dihydroxy-isovalerate [23]. To evaluate the essentiality of IlvY, we deleted *ilvY*(Δ*ilvY*) in two species of *Shigella, Shigella flexneri* 2a and *Shigella sonnei* and cultured wild-type and Δ*ilvY Shigella* in tryptic soy broth (TSB). IlvY deficient *Shigella* replicate at a similar rate and to a similar extent as wild-type *Shigella* (Fig. 2A-F) indicating that IlvY is not essential for bacterial replication under nutrient replete conditions. In contrast, growth of Δ*ilvY S. sonnei* and *S. flexneri* is reduced compared to wild-type *Shigella* in minimal broth lacking valine and isoleucine (Fig. 2G-L). Interestingly, Δ*ilvY S. sonnei* is unable to replicate in minimal broth whereas Δ*ilvY S. flexneri* is only modestly attenuated by the absence of IlvY (Fig. 2G-L).

**Figure 1.**
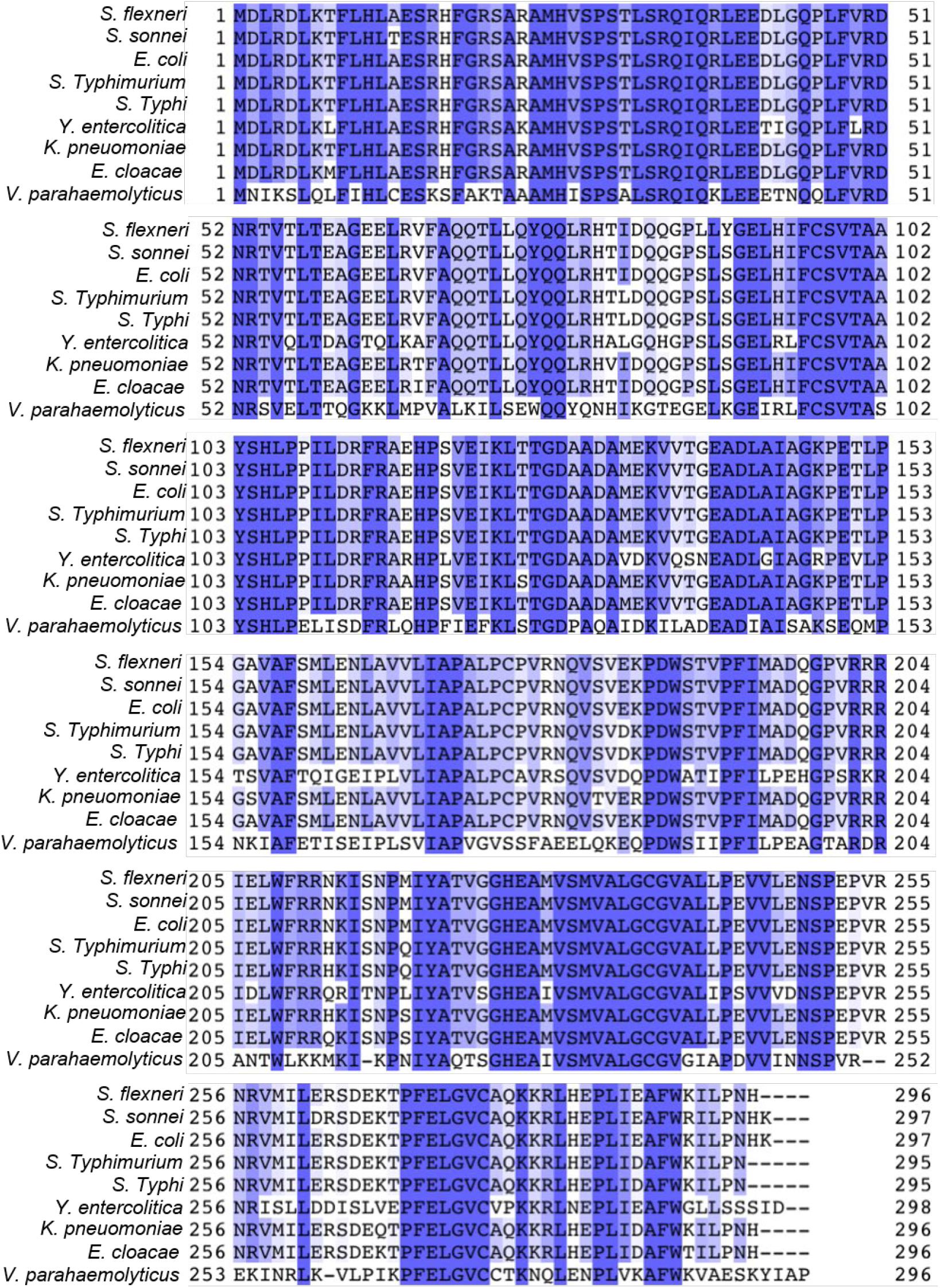
IlvY is highly conserved within Enterobacterales. Amino acid alignment of IlvY from *Shigella flexneri*, *Shigella sonnei*, *Escherichia coli, Salmonella enterica* serovar Typhimurium, *Salmonella enterica* serovar Typhi, *Yersinia entercolitica, Klebsiella pneumoniae, Enterobacter cloacae*, and *Vibrio parahemolyticus*. Coloring indicates percent identity across the sequences.

**Figure 2.**
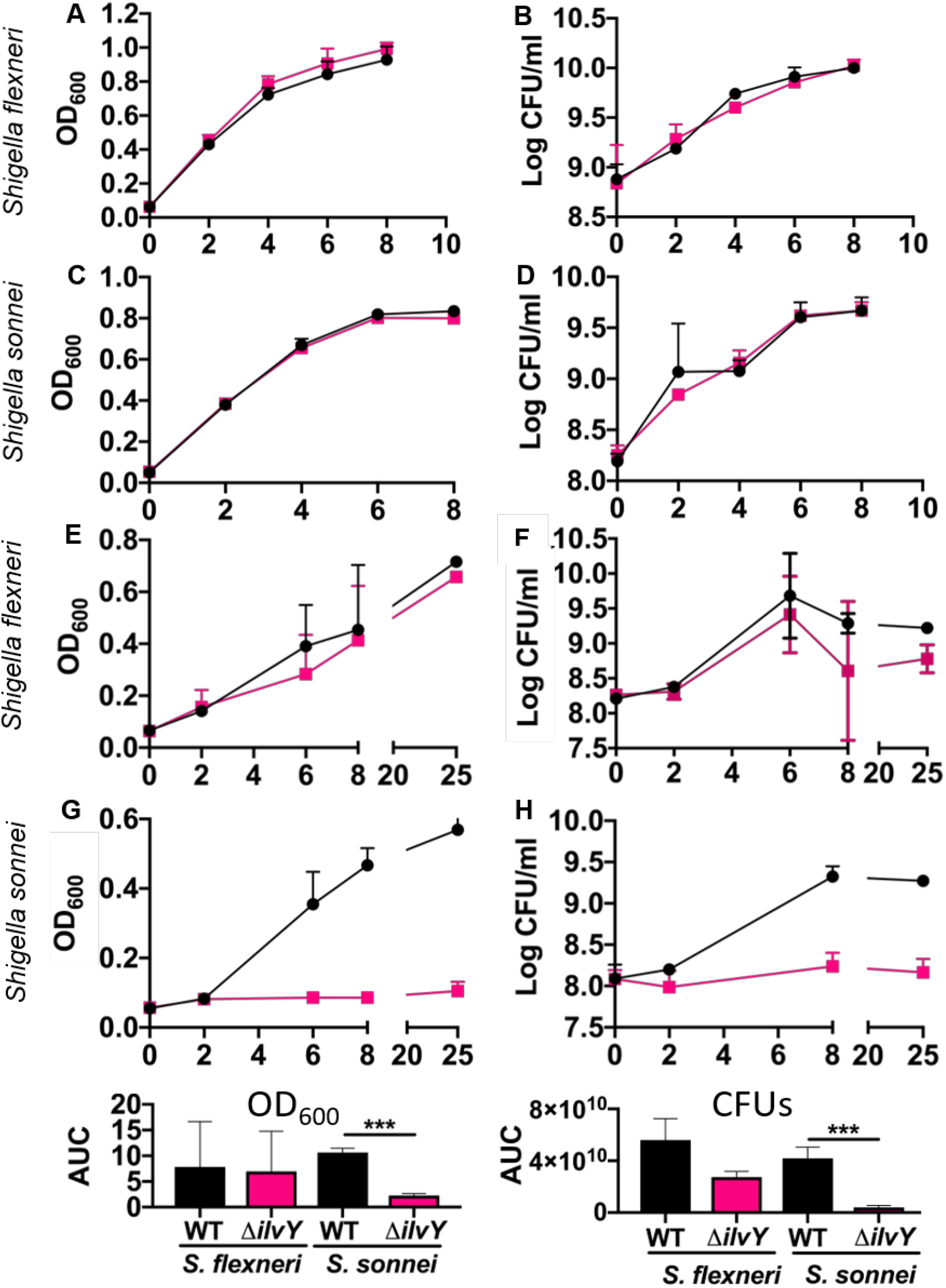
IlvY is dispensable during replication in rich broth, but important under nutrient limiting conditions. Replication of WT *S. flexneri* and WT *S. sonnei* (black) and ΔilvY *S. flexneri* and *S. sonnei* (pink) was quantified over time in tryptic soy broth (A-D) and minimal broth (E-H) by optical density 600 (A, C, E, G) and colony forming units (B, D, F, G). Lines connect mean values ± SD (n = 2 to 3) for each condition.

### *ilvC* transcription is activated under valine and isoleucine limiting conditions

Future efforts to develop therapeutics targeting IlvY will most likely require a high throughput assay to screen for compounds that impact IlvY activity. To that end we generated a transcriptional reporter construct to measure the activity of IlvY in *Shigella* in which the promoter for *ilvC* is driving expression of red fluorescent protein (RFP) (p*ilvC::*RFP). IlvY positively regulates *ilvC* transcription, thus RFP should be produced upon IlvY binding. To verify that IlvY does not activate *ilvC* transcription under all conditions, we cultured *Shigella* expressing constitutive RFP (pUltra-RFP), p*ilvC::*RFP, and promoter-less RFP (Δ::RFP) in TSB and measured RFP fluorescence over time. All *Shigella* reporter strains grew at similar rates in TSB indicating no differential effects from harboring the various expression constructs on replication (Fig. 3A and D). *Shigella* harboring Δ::RFP were used as controls for background autofluorescence. As expected, *Shigella* expressing constitutive RFP demonstrated increased RFP fluorescence over time. In contrast, the *Shigella* expressing p*ilvC::*RFP did not significantly increase in RFP fluorescence under these conditions (Fig. 3B and E). In contrast to growth in TSB, *Shigella* expressing p*ilvC::*RFP increased in fluorescence during growth in minimal broth (Fig. 3H and K). The strains expressing constitutive RFP are more fluorescent than the p*ilvC::*RFP reporter strains. Since the promoter driving RFP expression in the constitutive strain is optimized for maximal expression in Enterobacteriaceae, the higher fluorescence was not surprising [24].

**Figure 3.**
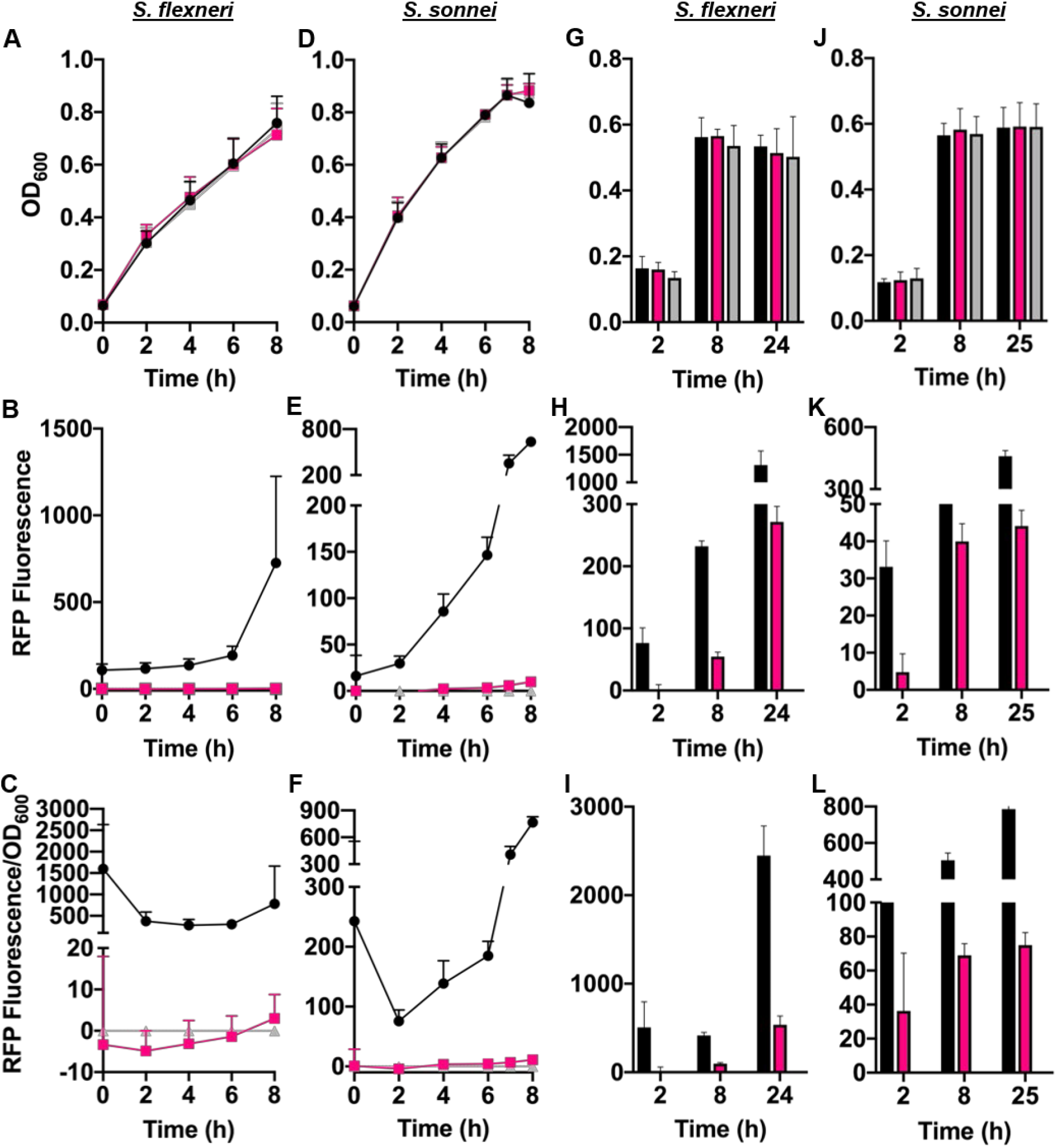
IlvY transcriptional reporters are activated *in vitro*. Replication (A, D, G, J) and RFP fluorescence (B, E, H, K) of *S. flexneri* and *S. sonnei* harboring pUltra-RFP (black), pUltra-p*ilvC::*RFP (pink), or pUltra-Δ::RFP (grey) was quantified over time in tryptic soy broth (A-F) and minimal broth (G-L). RFP fluorescence relative to OD_600_ was quantified for each strain (C, F, I, L). Lines connect mean values ±SD (n = 3 to 4) for each condition.

### *ilvC* expression is activated during infection of THP-1 cells

Previous transcriptional studies have shown that *ilvC* is upregulated upon infection [25]. Additionally, host cells initiate amino acid starvation following *Shigella* infection leading to reduced levels of isoleucine and leucine even in amino acid replete media [15]. Taken together, these data suggest that IlvC and by extension IlvY are likely important during intracellular replication. To determine whether *ilvC* transcription is activated during infection, we infected phorbol 12-myristate 13-acetate (PMA)-differentiated THP-1 cells with pUltra-RFP, p*ilvC::*RFP, and Δ::RFP expressing *Shigella*. RFP expressing bacteria were visible in THP-1 cells infected with p*ilvC::*RFP expressing *Shigella* (Fig. 4) suggesting that *ilvC* transcription is turned on in THP-1 cells. In concordance with results in Figure 3, *Shigella* containing pUltra-RFP were brighter than *Shigella* harboring p*ilvC::*RFP.

**Figure 4.**
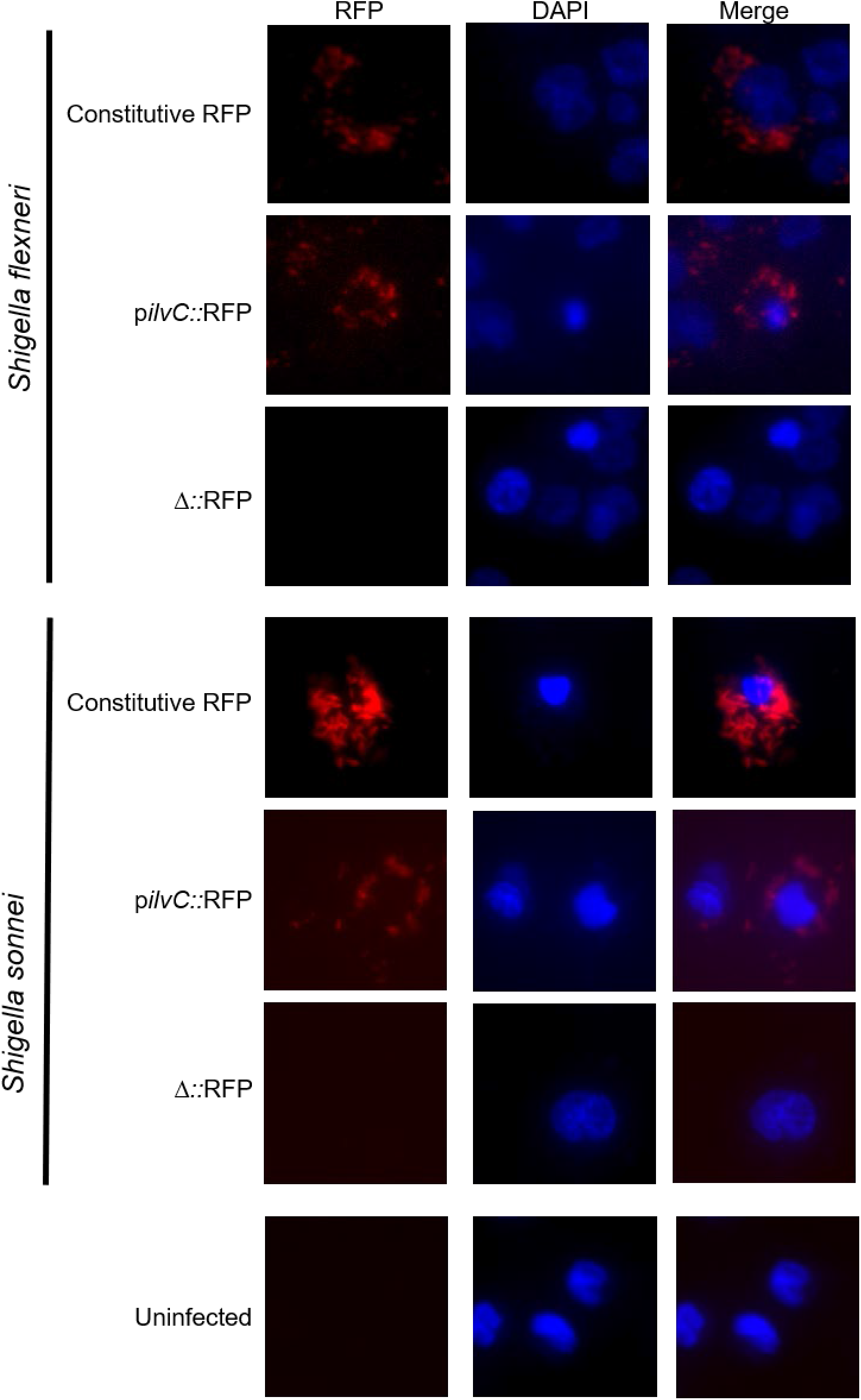
IlvY transcriptional reporters are activated during infection of THP-1 cells. THP-1 cells infected with *S. flexneri* and *S. sonnei* harboring pUltra-RFP, pUltra-p*ilvC::*RFP, or pUltra-Δ::RFP were imaged at 3 h post-infection. Cells were stained with Hoechst 33342 and imaged with a Nikon Eclipse T*i*.

Although the activation of the p*ilvC::*RFP reporter during infection of HCT-8 and THP-1 cells suggests that IlvY and IlvC are expressed, it is unclear whether IlvY is important for *Shigella* infection of human cells. We infected HCT-8 and THP-1 cells with WT and Δ*ilvY S. flexneri* and *S. sonnei*. WT and Δ*ilvY Shigella* did not show any difference in binding to or replication within HCT-8 cells (Fig. 5A-C). Unexpectedly, Δ*ilvY S. flexneri* was less efficient at binding to THP-1 cells than wild-type (Fig. 5D). While, this was not true for *S. sonnei*, ΔilvY *S. sonnei* was attenuated for intracellular replication at 4 hours post-infection (Fig. 5F). Taken together these data suggest that IlvY is involved during infection of human cells.

**Figure 5.**
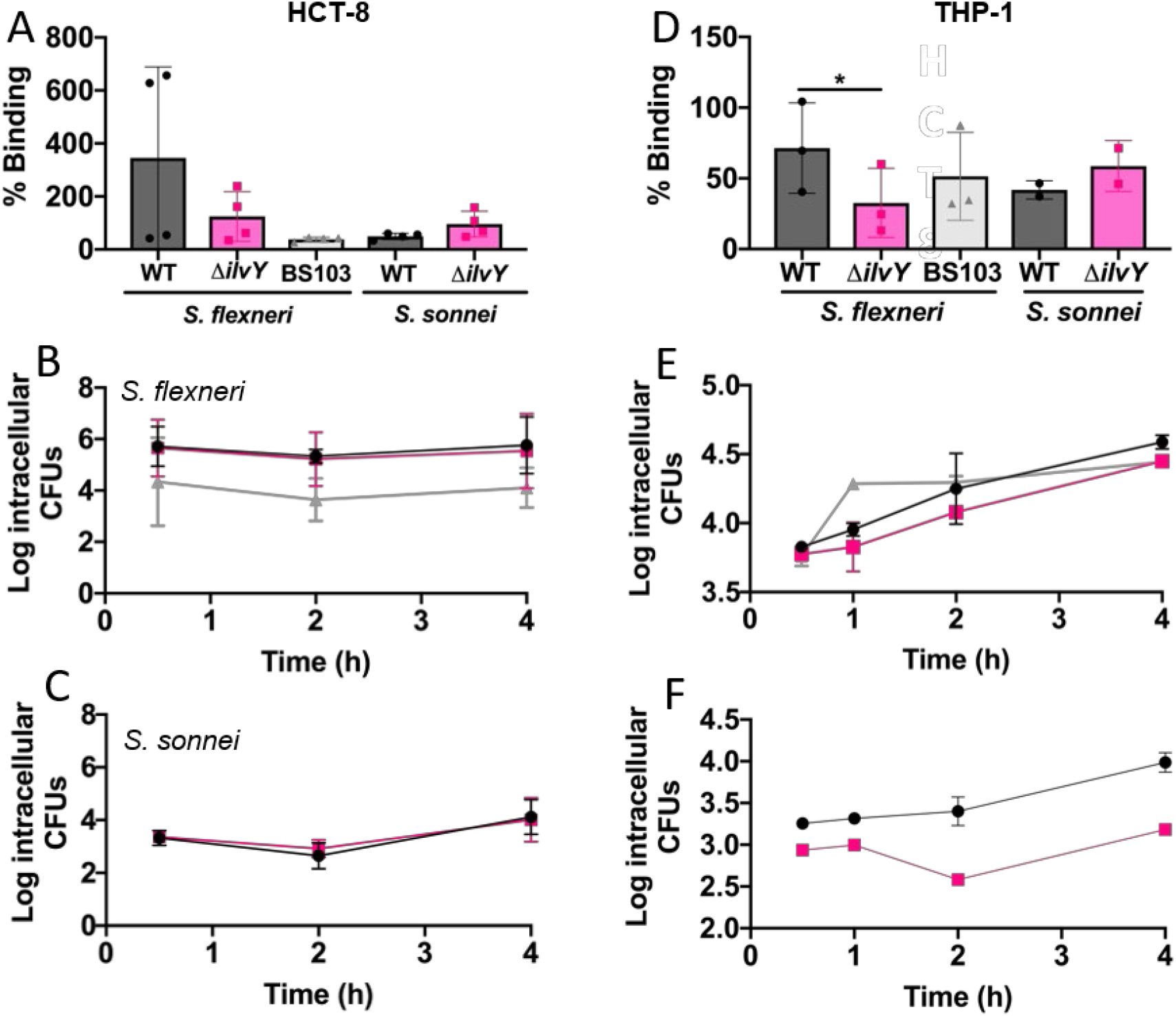
Binding and replication of *Shigella* in HCT-8 and THP-1 cells. Percent binding of *Shigella* strains to HCT-8 cells (A), and intracellular replication of *S. flexneri* (B) and *S. sonnei* (C) in HCT-8 cells. Percent binding of *Shigella* strains to THP-1 cells (D), and intracellular replication of *S. flexneri* (E) and *S. sonnei* (F) in THP-1 cells. Colors are consistent with Figure 2 with the exception of BS103 is grey. Lines connect mean values ± SD (*n =* 2) for each condition. One representative graph shown.

### In vivo study

Translating *Shigella in vitro* phenotypes to *in vivo* effects has been challenging due to the historic lack of mouse oral infection models. Current mouse models of oral infection require antibiotic pretreatment and/or diet modifications to induce susceptibility to infection. Based on the relatively recent discovery that *S. sonnei* has a Type VI Secretion System [26], we hypothesized that mice would be susceptible to orally administered *S. sonnei*. Therefore, we inoculated three mice orally with 5 x 10^5^ *S. sonnei* CFUs/mouse and monitored the time course of gastrointestinal infection (CFU/g tissue), a biomarker of inflammation (myeloperoxidase), and a biomarker of epithelial permeability (zonulin) (Fig. 6). While *S. sonnei* was not detected in the small intestine of inoculated mice, the large intestine of all mice had detectable *S. sonnei* at 24 hours (Fig. 6A). In addition, myeloperoxidase (a marker of neutrophil migration) was elevated at 24 hours and remained elevated at 72 hours (Fig. 6B). Furthermore, zonulin levels were reduced in infected mice suggesting that *S. sonnei* infection disrupted the gastrointestinal epithelium (Fig. 6C). Based on these results, the oral infection model was used to confirm the importance of IlvY for *S. sonnei in vivo* survival (Fig. 6D). We observed that the infection burden of mice infected with Δ*ilvY S. sonnei* was significantly less than mice infected with wild-type *S. sonnei* (Fig. 6D). While there was not a difference in the myeloperoxidase levels in the large intestine of mice infected with each strain, zonulin levels were significantly higher in mice infected with Δ*ilvY S. sonnei* (Fig. 6E-F). Taken together, these data indicate that *S. sonnei* lacking IlvY is attenuated for infection of mice, supporting our *in vitro* data.

**Figure 6.**
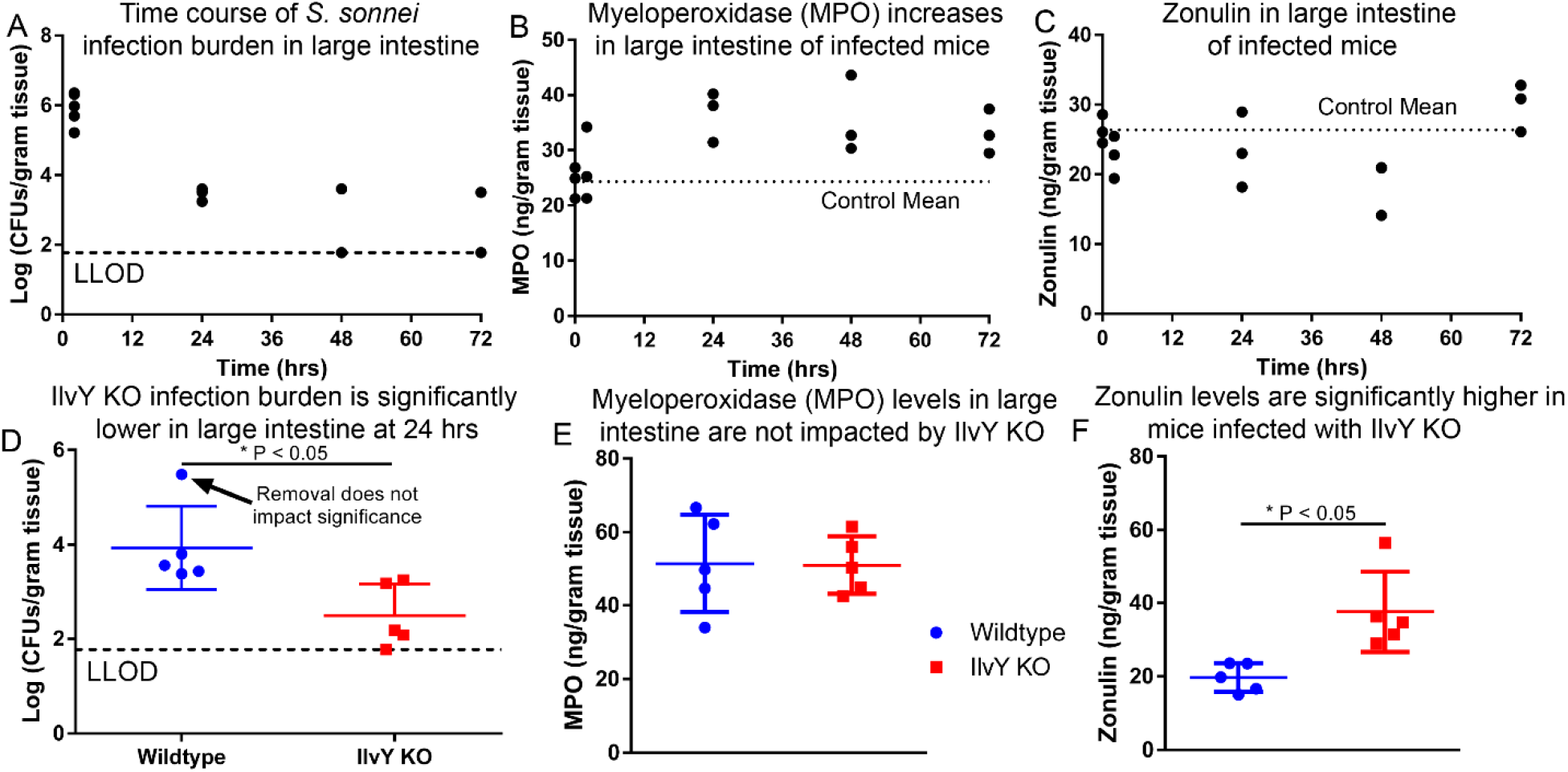
IlvY is important for *S. sonnei in vivo* survival. Wild-type C57Bl/6J mice were infected by oral gavage with 5 x 10^5^ CFU *S. sonnei* that were grown to log-phase in broth culture. At multiple time points post infection, gastrointestinal tissue was collected to characterize the infection burden (A). The observed infection in the large intestine was highest at 2 hours post infection, and all mice remained infected at 24 hours post infection. The inflammatory marker myeloperoxidase was significantly increased in the large intestine by 24 hours and remained elevated at 72 hours post infection (B). Zonulin, a marker of gastrointestinal epithelial permeability, was reduced over the first two days of infection but returned to baseline by 72 hours (C). To confirm whether IlvY has an important role in *S. sonnei in vivo* survival, Wild-type C57Bl/6J mice were infected by oral gavage with 5 x 10^8^ CFU wild-type *S. sonnei* or Δ*ilvY S. sonnei* that were grown to log-phase in broth culture. At 24 hours post infection, gastrointestinal tissue was collected to characterize the infection burden (D). Interestingly, mice infected with the Δ*ilvY S. sonnei* had a significantly lower infection burden compared to mice infected with wild-type *Shigella*. While the levels of myeloperoxidase were the same in both groups (E), zonulin levels were significantly higher in the large intestine of mice infected with Δ*ilvY S. sonnei* suggesting that intestinal permeability was not disrupted by infection with the IlvY deficient strain (F).

## Discussion

AMR is a global issue that is exacerbated by several factors including the overuse and misuse of antimicrobials, and roadblocks within the antibacterial development pipeline [5]. Since the development of antibiotics in the early 20th century, antibacterial therapeutics have targeted bacterial processes that are both distinct from mammalian pathways and essential for bacterial survival. The main cellular processes targeted by antibiotics include cell wall synthesis (β-lactams), protein synthesis (aminoglycosides and macrolides), DNA replication (fluoroquinolones), and folic acid synthesis (sulfonamides) [27]. Inhibition of any of these pathways culminates in bacterial cell death, thus placing significant selective pressure upon the bacteria to develop resistance to the antibiotics. Moreover, antibiotics are broadly acting and therefore target not only the pathogenic bacteria but also members of the microbiome. Antibacterial-induced alterations of the microbiome can lead to susceptibility to other bacterial infections such as *Clostridium difficile* [28]. Thus, it is important to identify novel targets for antibiotic development that neither inhibit essential cellular pathways nor decimate the microbiome.

Many pathogenic bacteria can replicate within host cells in addition to the extracellular environment, thus effective antibiotics must target bacteria in a variety of different niches within the human body. However, traditional antibiotics are bactericidal inside and outside of the host, which is conducive to the emergence of antibiotic resistance. Ideally, antibiotics would be effective at inhibiting infection within the host and inconsequential outside of the host. To achieve this, the pipeline of antibiotic target identification must be reworked to focus on processes that are important during infection and pathogenesis. Transcription factors are an ideal target for antibiotic development because they are important for establishment of infection and pathogenesis, but dispensable for survival. Targeting and inhibition of transcription factors during infection could ameliorate disease while not harming the microbiome or placing significant selective pressure on the pathogen. The concept of transcription factors as therapeutic targets has existed for a long time, but efficient targeting of has been difficult [29, 30]. This has largely been due to focusing primarily on developing compounds that occupy to the specific promoter to prevent transcription factor binding [8]. However, there are other domains/interfaces on transcription factors that could be targeted to modulate gene expression [30]. Availability of targeting sites varies among all transcription factors, but sites that represent possible targets include inducer or inhibitor binding sites, dimerization domains, interfaces important for interaction with other transcriptional machinery, or clefts involved in conformational changes to allow for binding and/or transcriptional activation/repression after DNA binding. Recent studies involving anticancer and antimicrobial drugs targeting transcription factors have shown promise [9, 11, 31]. With respect to targeting transcription factors in *Shigella*, researchers have primarily focused on VirF, the master regulator of *Shigella* virulence factor expression [11, 12]. Much of VirF regulation of virulence factors is mediated through activation of a second transcription factor VirB [32]. It is important to note that *virB* expression can be activated independently of VirF [33], thus targeting VirF to inhibit virulence gene expression is likely to be leaky. Moreover, VirF is not highly conserved across different genera of bacteria, limiting the usefulness of a small molecule that targets VirF. Therefore, targeting a transcription factor like IlvY may provide a promising alternative since it is highly conserved across the order of Enterobacterales (Fig. 1) and tightly controls transcription of its two targets [22].

Another factor to consider when identifying potential therapeutic targets is the conservation amongst bacteria. Previous approaches to targeting pathogenic bacteria have focused primarily on developing compounds that inhibit Type III Secretion Systems. However, this method would require development of compounds that would target important virulence factors for each pathogenic bacterial species and the targets of many of these inhibitors have not been identified [34]. By targeting conserved transcription factors the drugs will be effective against a wide swath of enteric bacteria that cause disease in humans. Importantly, since the transcription factors are non-essential, inhibition of a transcription factor expressed by a constituent of the microbiome is unlikely to negatively impact the survival.

*Shigella* is a significant global pathogen with increasing AMR that is closely related to other enteric pathogens including enteroinvasive *Escherichia coli, Salmonella enterica*, and *Yersinia entercolitica*. Thus, it is a model organism for identifying and elucidating the importance of transcription factors during both *in vitro* and *in vivo* infection *Shigella* exhibits a complex infection process as it replicates efficiently in the extracellular milieu (intestinal lumen and interstitial space in the lamina propria) and intracellularly (intestinal epithelial cells and macrophages). Our work demonstrates that the highly conserved, non-essential transcription factor, IlvY, is a potential target for antimicrobial drug development (Fig. 1). IlvY regulates two genes *ilvC*, which encodes a ketol-acid reductoisomerase, and *ilvY*[22]. IlvC catalyzes the conversion of acetohydroxybutyrate or acetolactate to 2,3-dihydroxy-isovalerate as part of the valine and isoleucine biosynthetic pathways. Interestingly, *ilvC* and *ilvY* share a promoter region but are oriented in opposite directions [22, 23]. Notably, IlvY activity is modulated by the substrates of IlvC [35]. Upon binding of acetohydroxybutyrate or acetolactate to IlvY, *ilvC* transcription is activated while *ilvY* transcription is repressed. It has been shown previously that *ilvY* is upregulated during intracellular replication [25], indicating that it is involved in intracellular growth. Moreover, cells initiate amino acid starvation following *Shigella* infection that results in reduced intracellular levels of isoleucine and leucine even though the cells are cultured in the presence of all essential amino acids [15]. Thus, IlvY mediated activation of *ilvC* is likely important for survival under amino acid starvation when isoleucine and valine are less available.

To assess the importance of IlvY, the *ilvY* gene was knocked out of two *Shigella* species, *S. flexneri* 2a and *S. sonnei*, which together cause the majority of cases worldwide [36]. For development of antimicrobial compounds that do not kill bacterial cells, it is critical that the targeted transcription factor is non-essential. As expected, deletion of *ilvY* in either *S. flexneri* 2a or *S. sonnei* did not result in significant differences in growth rates in a rich medium, tryptic soy broth (Fig. 2A-D). Although *ilvY* deleted strains grew well in rich broth with sufficient quantities of nutrients, Δ*ilvY S. flexneri* 2a and Δ*ilvY S. sonnei* were attenuated compared to wild-type strains in minimal broth lacking added valine or isoleucine (Fig. 2E-H). Interestingly, Δ*ilvY S. sonnei* did not replicate in minimal broth whereas Δ*ilvY S. flexneri* 2a replication was only modestly reduced in minimal broth. It is unclear why the loss of IlvY dramatically effects only one of the two *Shigella* species. To effectively identify a promising drug candidate, it is useful to have a robust assay to monitor transcription factor activity. Since the promoter region of *ilvY* and *ilvC* has been extensively studied for being a prototypical LysR type regulon, the bounds of the promoter and critical binding sites are known [22, 23]. Given that the promoter region is well defined, it is feasible to generate a transcriptional reporter construct to measure regulatory activity of IlvY. We generated a transcriptional reporter construct in which the promoter for *ilvC* is driving expression of RFP. Upon activation of *ilvC* gene expression, RFP should be expressed within the cell. Growth of *S. flexneri pilvC::RFP* and *S. sonnei pilvC::RFP* reporter strains in minimal broth without exogenous valine or isoleucine resulted in increased RFP expression (Fig. 3G-L), demonstrating the utility of this reporter for measuring IlvY activity. Importantly, the p*ilvC*::RFP reporters did not turn on in TSB (Fig. 3A-F) indicating that RFP expression is specific to conditions conducive to *ilvC* transcription.

Although minimal broth is sufficient to induce *ilvC* transcriptional activation, it is not representative of the physiologic environment during infection. An effective drug target must be activated during infection of the host. During infection of HCT-8 cells and THP-1 cells, intracellular *S. flexneri pilvC::RFP* and *S. sonnei pilvC::RFP* reporter strains expressed RFP (Fig. 4), indicating that *ilvC* expression is activated by IlvY during infection. Activation of the p*ilvC*::RFP construct during infection suggests that IlvC important for intracellular replication. However, it does not necessarily mean that IlvC is indispensable for replication within host cells. New putative therapeutic targets for treating *Shigella* infection should be able to inhibit intracellular replication, thus the transcription factor needs to be involved in mediating efficient intracellular replication. Deletion of *ilvY* in both *S. flexneri* and *S. sonnei* attenuated intracellular replication in THP-1 cells at 4.5 hours post-infection compared to WT (Fig. 5). These data suggest that IlvY is an important regulator of intracellular replication in human cells. Since IlvY does not directly regulate an essential gene or classical virulence factors, IlvY is likely exerting its effect through modulation of biochemical pathways and response to host derived antimicrobial agents. It has been shown previously that *Shigella* infection induces a 4 hour period of amino acid stress in infected host cells leading to reduced intracellular levels of branched chain amino acids [14–16]. As such *Shigella* would need to generate its own valine and isoleucine to survive within the amino acid deprived cytosol. Inhibition of the valine and isoleucine biosynthetic pathway either genetically (Δ*ilvY*) or chemically would therefore lead to attenuated replication in the cytosol. Additionally, IlvY has been connected to the nitrosative stress response [14]. IlvD, a dihydroxy-acid dehydratase, is directly downstream of IlvC in the valine and isoleucine biosynthetic pathways, and contains an iron-sulfur cluster, which can be damaged by host-derived nitrosative stress. Inactivation of IlvD by nitrosative stress will lead to a build of upstream products (e.g., substrates for IlvC) and activation of IlvY. Taken together IlvY is an optimal target for developing therapeutics.

A significant hindrance to the understanding of *Shigella* pathogenesis has been the lack of a suitable animal model for many years. Researchers have commonly used guinea pigs and rabbit ileal loops to model *Shigella* infection in animals [26, 37–39]. However, given the route of inoculation or ex vivo nature of these models, they do not mimic the route of infection in humans. For the first time, we have developed an oral adult mouse model of *Shigella* infection model that does not require antibiotic pretreatment or special diet modifications. Why mice exhibit differences in susceptibility to orally administered *S. flexneri* and *S. sonnei* is unclear, but it may be due to *S. sonnei* having a Type VI Secretion System [26]. *S. sonnei* is detectable in the cecum/large intestine up to 48h post-infection (Fig. 6A). Interestingly, myeloperoxidase increased over time in infected mice indicating recruitment of neutrophils to the intestines (Fig. 6B). In addition, the reduction in zonulin suggests that the epithelial permeability is disrupted by infection (Fig. 6C). To evaluate the impact of IlvY on *S. sonnei in vivo* infection, mice were infected with either WT or Δ*ilvY S. sonnei* and sacrificed at 24 h post-infection. Mice infected with Δ*ilvY S. sonnei* contained fewer CFUs in their cecum and large intestine at 24 h post-infection compared to mice infected with WT *S. sonnei* (Fig. 6D). While myeloperoxidase levels were not impacted by *ilvY* deletion, zonulin levels were significantly higher in mice infected with Δ*ilvY S. sonnei* (Fig. 6F). This difference may be related to our observation that Δ*ilvY S. sonnei* replication was reduced in THP-1 cells (macrophage like cells) but not HCT-8 cells (intestinal epithelial cells). If *ilvY* deletion diminishes *Shigella* macrophage invasion and subsequent pyroptosis, it is possible that this relative reduction in macrophage pyroptosis does not lead to the resulting zonulin decrease observed with wild-type *Shigella* infection. Overall, these data demonstrate that this model of *Shigella* oral infection is suitable for evaluating the roles of different genes during infection.

*Shigella* transcription factors represent a new target for therapeutic development. IlvY is a candidate transcription factor for antibacterial drug development because it is non-essential, conserved across enteric pathogens, and it is important for infection.

## Materials and Methods

### Cell lines

Human ileocecal adenocarcinoma HCT-8 cells (ATCC) and THP-1 cells (ATCC) were maintained in Advanced RPMI 1640 (ThermoFisher Scientific) supplemented with 10% fetal bovine serum and 1X penicillin-streptomycin (ThermoFisher Scientific) (complete media).

### Bacterial strains

*Shigella flexneri* 2a 2457T (BEI resources) and *Shigella sonnei* WRAIR (BEI resources) were used for all experiments.

**Table 1.** Strains

| Table 1. Strains |  |
| --- | --- |
| Strain | Source |
| *S. flexneri* 2a 2457T | BEI Resources |
| *S. sonnei* WRAIR | BEI Resources |
| *S. flexneri* 2a 2457T pUltra-mRFP1 | This paper |
| *S. flexneri* 2a 2457T pUltra-*pilvC*::mRFP1 | This paper |
| *S. flexneri* 2a 2457T pUltra-Δ::mRFP1 | This paper |
| *S. flexneri* 2a 2457T pUltra-mRFP1 pUC19-GFP | This paper |
| *S. flexneri* 2a 2457T pUltra-*pilvC*::mRFP1 pUC19-GFP | This paper |
| *S. flexneri* 2a 2457T pUltra-Δ::mRFP1 pUC19-GFP | This paper |
| BS103 | Helen Wing |
| *S. sonnei* WRAIR pUltra-mRFP1 | This paper |
| *S. sonnei* WRAIR pUltra-*pilvC*::mRFP1 | This paper |
| *S. sonnei* WRAIR pUltra-Δ::mRFP1 | This paper |
| *S. sonnei* WRAIR pUltra-mRFP1 pUC19-GFP | This paper |
| *S. sonnei* WRAIR pUltra-*pilvC*::mRFP1 pUC19-GFP | This paper |
| *S. sonnei* WRAIR Kan<sup>R</sup> | This paper |
| *S. sonnei* WRAIR Kan<sup>R</sup> Δ*ilvY* | This paper |

**Table 2.** Plasmids

| Table 2. Plasmids |  |  |
| --- | --- | --- |
| Plasmid | Resistance | Source |
| pGRG25 | Ampicillin | [40] |
| pGRG25-Kan<sup>R</sup> | Ampicillin | This paper |
| pUltra-mRFP1 | Kanamycin | [24] |
| pTOX3 | Chloramphenicol | [41] |
| pUltra-*pilvC*::mRFP1 | Kanamycin | This paper |
| pUltra-Δ::mRFP1 | Kanamycin | This paper |
| pTOX3 Δ*ilvY* | Chloramphenicol | This paper |
| pUC19-GFP | Ampicillin | This paper |
| pUltra-*ilvY* | Kanamycin | This paper |
| pBAD18 | Ampicillin | [42] |
| pBAD18-*ilvY* | Ampicillin | This Paper |

### Fluorescent and complement strain generation

pUltra-RFP was purified using ZymoPure II Plasmid Midi Prep kit (Zymo Research), digested with BamHI and EcoRI, and gel purified. The *ilvC* promoter was synthesized as a gBlock (Table 4, GeneWiz) and mRFP1 was PCR amplified using primers listed in Table 3 and purified using DNA Clean and Concentrator-5 kit (Zymo Research). pUC19 was digested with PstI and pBAD18 was digested with EcoRI and HindIII, and then gel purified. *eGFP* and *ilvY* were amplified using primers listed in Table 3 and purified as described above. Constructs were assembled using the Gibson assembly master mix (NEB) according to manufacturer’s directions and transformed into NEB5α competent cells (NEB). Correct constructs were electroporated into *Shigella* using Amaxa 96-well shuttle system (Lonza).

**Table 3.**
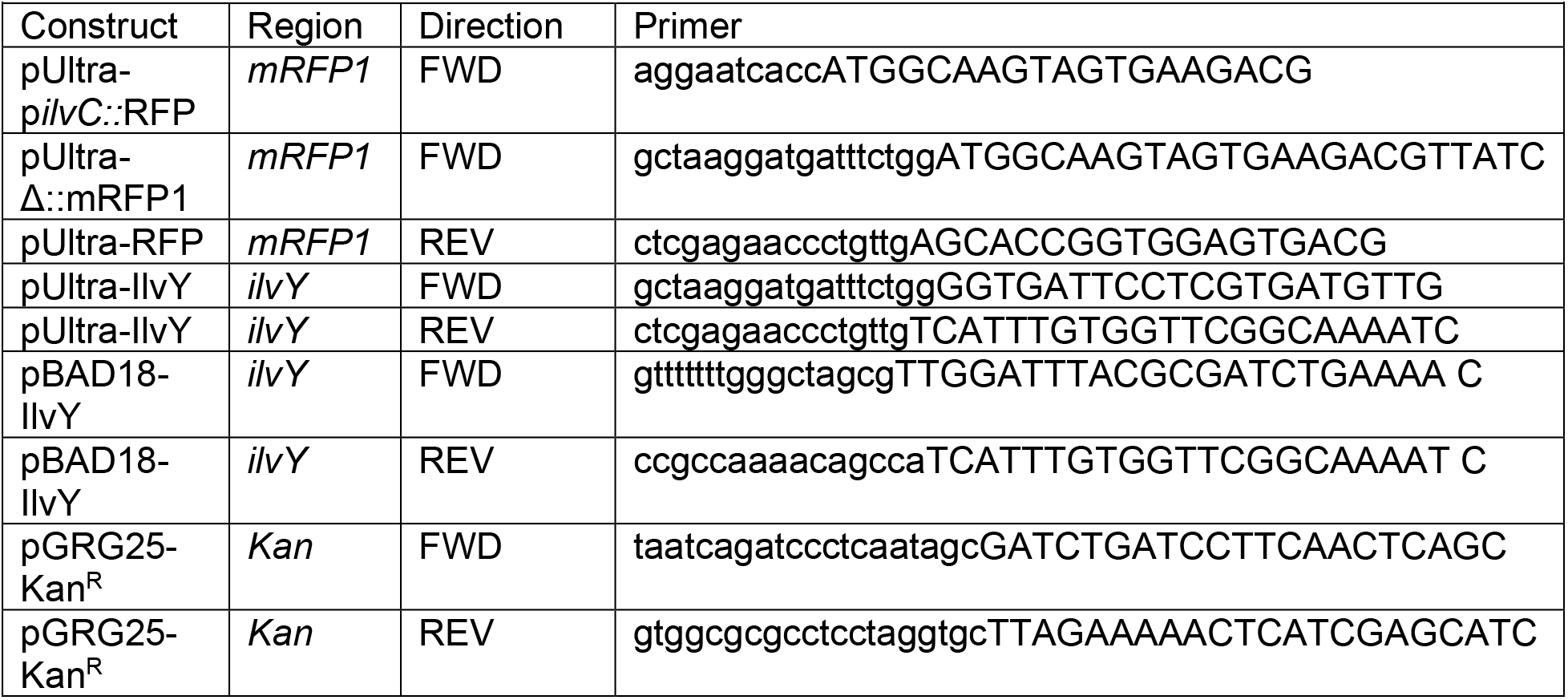
Primers

**Table 4.**
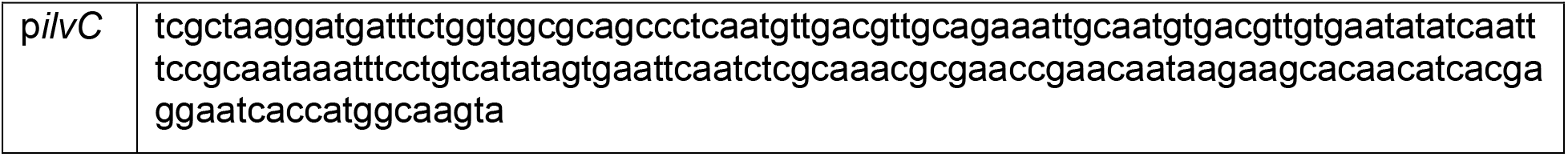
gBlocks

### Mutant generation

500 bp upstream and 500 bp downstream regions of each gene were amplified from *S. flexneri* 2a 2457T genomic DNA. The 500 bp fragments were inserted into SmaI digested pTOX3 plasmid using the Gibson assembly master mix. Correct pTOX3 constructs were transformed into SM10λpir *E. coli*. pTOX3 constructs were conjugated into *Shigella* using SM10λpir at 37°C. *Shigella* merodiploids were isolated on minimal agar containing 1X M9 salts, 0.5 mM MgSO_4_, 0.1 mM CaCl_2_, 1% glucose, 20 μg/ml tryptophan, 12.5 μg/ml nicotinic acid, 45 μg/ml methionine, 25 μM Fe(III)Cl3 in 50 μM citric acid, 20 μg/ml chloramphenicol, and 3% Bacto agar. Potential *Shigella* merodiploids were then struck out on MacConkey agar with 20 μg/ml chloramphenicol to differentiate from *E. coli*. Lactose fermentation negative, chloramphenicol resistant colonies were then inoculated into 2 ml LB broth with 2% glucose and incubate for 2 h at 37°C. Bacteria were washed twice in 1X M9 salts with 2% rhamnose and plated on to rhamnose agar: 1X M9 salts, 0.5 mM MgSO_4_, 0.1 mM CaCl_2_, 0.2% rhamnose, 20 μg/ml tryptophan, 0.2% casamino acids, 12.5 μg/ml nicotinic acid, 25 μM Fe(III)Cl3 in 50 μM citric acid, and 3% Bacto agar. Unless otherwise stated, plates were incubated at 30°C. Mutants were identified using colony PCR.

### In vitro infection assays

HCT-8 cells were seeded at 2.5×10^4^ cells per well in a 96-well plate 3 days prior to infection. 1 day prior to infection, the media on the HCT-8 cells was removed and replaced with antibiotic free complete media. THP-1 cells were seeded at 8×10^4^ cells per well in complete media supplemented with 100 nM phorbol myristate acetate (PMA) in 96-well plate 3 days prior to infection. After 48 h the PMA containing media was removed and replaced with antibiotic free complete media lacking PMA. Overnight cultures of *S. flexneri* and *S. sonnei* in TSB (ThermoFisher Scientific) containing 0.1% deoxycholate (ThermoFisher Scientific) were diluted 50-fold in TSB supplemented with 0.1% deoxycholate and incubated at 37°C 225 RPM for 2.5h. Bacteria were washed in PBS and resuspended in complete media without antibiotics. HCT-8 and THP-1 cells were infected at a multiplicity of infection of 100 and 10, respectively. *Shigella* were centrifuged on to the cells at 1000 × g for 10 min at ambient temperature, then incubated at 37°C with 5% CO_2_ for an additional 30 min. The inoculum was removed, and the cells were washed 3 times with PBS. To quantify percent *Shigella* binding to cells, cells were lysed following PBS washes in 0.1% triton-X 100 (Sigma-Aldrich) in PBS and plated on to LB agar. To quantify intracellular replication complete media supplemented with 20 ug/ml gentamicin was added to the cells for 30 min at 37°C with 5% CO_2_ to kill extracellular bacteria.

Following gentamicin treatment, cells were washed one time with PBS then replaced with antibiotic free complete media. At 30 min, 1, 2, and 4 h post-binding cells were lysed, serially diluted, and plated for colony forming units (CFUs) on LB agar.

### Fluorescence microscopy

Infections were performed as described above with the following modifications. Cells were cultured in a black wall 96-well plates (Corning) to reduce background fluorescence. After removal of the inoculum cells were incubated at 37°C with 5% CO_2_ for an additional 3 h. 15 min prior to visualization on the microscope, media was removed, cells were washed 3 times with PBS, and phenol red free RPMI 1640 supplemented with L-glutamine and 10% FBS was added to the wells. To visualize the nuclei, 100 ug/ml Hoechst 33342 was added to each well. Images were acquired using a Nikon Eclipse Ti with a Hamamatsu digital camera C10600 camera (Orca-R^2^) and Volocity software (Quorum Technologies).

### Viability growth curves

Overnight cultures of *S. flexneri* and *S. sonnei* were back diluted to OD_600_ of 0.05 in appropriate growth media. OD_600_ was measured at indicated time points by transferring 100 μl aliquots of the cultures to clear 96-well plates and measured on a BioTek ELx600. Serial dilutions of cultures were plated on to LB agar at indicated time points to quantify the number of CFUs.

### Fluorescence growth curves

Overnight cultures of *S. flexneri* and *S. sonnei* transcriptional reporter strains were back diluted to OD_600_ of 0.05 in appropriate growth media. To measure OD_600_ and RFP fluorescence 100 μl aliquots of the cultures were added to 96-well black-walled plates and measured using BioTek ELx600 and FLx600. For growth curves performed in minimal broth, bacteria were washed 2 times in PBS prior to resuspension in minimal broth. Minimal broth contained 1X M9 salts, 0.5 mM MgSO_4_, 0.1 mM CaCl_2_, 1% glucose, 20 μg/ml tryptophan, 12.5 μg/ml nicotinic acid, 45 μg/ml methionine, 25 μM Fe(III)Cl3 in 50 μM citric acid, and 50 μg/ml kanamycin.

### Mouse infections

WT and Δ*ilvY S. sonnei* strains harboring pUltra-RFP and pUC19-GFP were subcultured in TSB containing 0.1% sodium deoxycholate, 50 μg/ml ampicillin, and 50 μg/ml kanamycin at 37°C, shaking at 225 rpm for 3 h. *S. sonnei* was washed and resuspended in PBS at 2.5 x 10^9^ CFU/mL. WT C57BL/6J mice (The Jackson Laboratory) were orally gavaged with 5 x 10^8^ CFUs. At 2 and 24 h post-infection mice were sacrificed, small intestinal tissue and colonic tissues were harvested separately, and flash frozen in liquid nitrogen. To quantify CFUs in intestinal tissue samples, tissues were homogenzied in 3 times the volume of PBS using a handheld homogenizer, and plated on to LB agar containing 50 μg/ml kanamycin and 50 μg/ml ampicillin.

### ELISAs

Intestinal tissue samples were homogenized in 3 times the volume of PBS using a handheld homogenizer. Samples were aliquoted in 1 mL quantities and flash frozen in liquid nitrogen and stored at −80°C until ready for use. Aliquoted samples, assay reagents and materials were allowed to warm to room temperature for 30 minutes. Samples were centrifuged at 3000 rpm for 20 minutes and the supernatants were transferred to the ELISA plate in duplicate. Assays were performed for each of the following proteins according to the manufacturer’s instructions: Myeloperoxidase (MyBioSource), Alkaline Phosphatase (MyBioSource), Zonulin (MyBioSource), C-reactive protein (MyBioSource). The optical density was read at 450 nm on an EnVision plate reader (PerkinElmer). Between assays, samples were stored at 4°C and used within 5 days.

### Animal ethics statement

All animal procedures and protocols were approved by the institutional animal care and use (IACUC) committee at the University of Washington (protocol Number: 2154-01) and all efforts were made to minimize animal discomfort and suffering. The University of Washington is accredited by the Association for the Assessment and Accreditation of Laboratory Animal Care, International (AAALAC). The Office of Laboratory Animal Welfare of the National Institutes of Health (NIH) has approved the University of Washington (#A3464-01), and this study was carried out in strict compliance with the Public Health Service (PHS) Policy on Humane Care and Use of Laboratory Animals.

### Statistics

GraphPad Prism software (GraphPad, La Jolla, CA) was used for statistical analysis. For determining linear relationships, linear regression analysis was performed. For *in vitro* and *in vivo* experiments, bacterial counts and biomarkers in the cells/intestines were compared using unpaired *t*-tests. Data were considered statistically significant for *P*≤ 0.05.

## Acknowledgements

We would like to thank Professor Helen Wing for providing *Shigella flexneri* BS103 and Raiven Yoes for their discussions about the work. This work was funded by Tres Cantos Open Lab Foundation (Grant: TC264).

## Contributions

S.A., M.K.H., M.A., A.S., B.M. designed the experiments. M.K.H. and L.R. conducted the molecular biology included. S.A., M.K.H., and L.R. analyzed and interpreted the data. S.A. and M.K.H. wrote the manuscript.

## Additional Information

Correspondence and request for materials should be addressed to S.A.

